# Genetic correlations among psychiatric and immune-related phenotypes based on genome-wide association data

**DOI:** 10.1101/070730

**Authors:** Daniel S. Tylee, Jiayin Sun, Jonathan L. Hess, Muhammad A. Tahir, Esha Sharma, Rainer Malik, Bradford B. Worrall, Andrew J. Levine, Jeremy J. Martinson, Sergey Nejentsev, Doug Speed, Annegret Fischer, Eric Mick, Brian R. Walker, Andrew Crawford, Struan F.A. Grant, Constantin Polychronakos, Jonathan P. Bradfield, Patrick M. A. Sleiman, Hakon Hakonarson, Eva Ellinghaus, James T. Elder, Lam C. Tsoi, Richard C. Trembath, Jonathan N. Barker, Andre Franke, Abbas Dehghan, The 23andMe Research Team, The Inflammation Working Group of the CHARGE Consortium, The METASTROKE Consortium of the International Stroke Genetics Consortium, The Netherlands Twin Registry, The neuroCHARGE Working Group, The Eating Disorders Working Groups of the Psychiatric Genomics Consortium, The Obsessive Compulsive Disorder and Tourette Syndrome Working Group, Stephen V. Faraone, Stephen J. Glatt

**Affiliations:** Psychiatric Genetic Epidemiology & Neurobiology Laboratory (PsychGENe Lab); Departments of Psychiatry and Behavioral Sciences & Neuroscience and Physiology; SUNY Upstate Medical University; Syracuse, NY, U.S.A; Institute for Stroke and Dementia Research, Klinikum der Universität München, Ludwig-Maximilians-University (LMU), Munich, Germany; Departments of Neurology and Public Health Sciences, University of Virginia School of Medicine, Charlottesville, VA, U.S.A; Department of Neurology, David Geffen School of Medicine, University of California Los Angeles, Los Angeles, CA, U.S.A.; Department of Infectious Diseases and Microbiology, Graduate School of Public Health, University of Pittsburgh, PA, U.S.A.; Department of Medicine, University of Cambridge, Cambridge, U.K.; Aarhus Institute for Advanced Studies and University College London, London, U.K.; UCL Genetics Instutute, University College London, United Kingdom WC1E 6BT; Institute of Clinical Molecular Biology, Christian Albrechts University of Kiel, Kiel, Germany; Department of Quantitative Health Sciences, University of Massachusetts Medical School, Worcester, MA, U.S.A.; BHF Centre for Cardiovascular Science, Queen’s Medical Research Institute, University of Edinburgh, Edinburgh, EH16 4TJ, U.K.; School of Social and Community Medicine, MRC Integrated Epidemiology Unit, University of Bristol, Bristol, BS8 2BN, UK; Center for Applied Genomics, Division of Human Genetics, The Children’s Hospital of Philadelphia, Philadelphia, PA, U.S.A.; Division of Endocrinology and Diabetes, The Children’s Hospital of Philadelphia, Philadelphia, PA, U.S.A.; Department of Pediatrics, Perelman School of Medicine, University of Pennsylvania, Philadelphia, PA, U.S.A.; Institute of Diabetes, Obesity and Metabolism, Perelman School of Medicine, University of Pennsylvania, Philadelphia, PA, U.S.A.; Endocrine Genetics Laboratory, Department of Pediatrics and the Child Health Program of the Research Institute, McGill University Health Centre, Montreal, Quebec, Canada; Quantinuum Research LLC, San Diego, CA, U.S.A.; Department of Dermatology, Veterans Affairs Hospital, University of Michigan, Ann Arbor, Michigan, United States of America; Department of Biostatistics, University of Michigan, Ann Arbor, Michigan, United States of America; Division of Genetics and Molecular Medicine, King’s College London, London, UK; Department of Biostatistics and Epidemiology, MRC-PHE Centre for Environment and Health, School of Public Health, Imperial College London; 23andMe, Inc., Mountain View, CA, USA; K.G. Jebsen Centre for Research on Neuropsychiatric Disorders, University of Bergen, Bergen, Norway

**Author notes:** To whom correspondence should be addressed: SUNY Upstate Medical University, 750 East Adams Street, Syracuse, NY 13210.

**Keywords:** allergy, anorexia nervosa, attention deficit-hyperactivity disorder, autoimmune disorder, bipolar disorder, celiac disease, childhood ear infection, C-reactive protein, Crohn’s disease, genetic correlation, genome-wide association, hypothyroidism, major depression, neuroticism, obsessive schizophrenia, primary biliary cirrhosis, rheumatoid arthritis, smoking, systemic lupus erythematosus, Tourette syndrome, tuberculosis susceptibility, type 1 diabetes, ulcerative colitis

## Abstract

Individuals with psychiatric disorders have elevated rates of autoimmune comorbidity and altered immune signaling. It is unclear whether these altered immunological states have a shared genetic basis with those psychiatric disorders. The present study sought to use existing summary-level data from previous genome-wide association studies (GWASs) to determine if commonly varying single nucleotide polymorphisms (SNPs) are shared between psychiatric and immune-related phenotypes. We estimated heritability and examined pair-wise genetic correlations using the linkage disequilibrium score regression (LDSC) and heritability estimation from summary statistics (HESS) methods. Using LDSC, we observed significant genetic correlations between immune-related disorders and several psychiatric disorders, including anorexia nervosa, attention deficit-hyperactivity disorder, bipolar disorder, major depression, obsessive compulsive disorder, schizophrenia, smoking behavior, and Tourette syndrome. Loci significantly mediating genetic correlations were identified for schizophrenia when analytically paired with Crohn’s disease, primary biliary cirrhosis, systemic lupus erythematosus, and ulcerative colitis. We report significantly correlated loci and highlight those containing genome-wide associations and candidate genes for respective disorders. We also used the LDSC method to characterize genetic correlations amongst the immune-related phenotypes. We discuss our findings in the context of relevant genetic and epidemiological literature, as well as the limitations and caveats of the study.

## Introduction

The biological bases of major psychiatric disorders have been studied for decades, yet they remain largely unresolved. Evidence from both clinical and biomedical literature has demonstrated that individuals with these conditions show differences in circulating immunologic markers, functional capacities of isolated immune cells, and atypical prevalence of clinical immune-related phenotypes compared to individuals not affected by psychiatric or neurodevelopmental disorders.^1–10^ It remains unclear what roles (if any) altered immunologic functions may play in the major psychiatric phenotypes, though plausible mechanisms linking altered immune functions with neurobiological changes during early brain development and in fully developed adults have been identified.^11–18^ While some studies have already suggested potential genetic bases for the immune dysregulation observed in a subset of psychiatric patients,^19–22^ the extent to which co-occurrence or segregation of clinical phenotypes may be influenced by similarities in genome-wide genetic risk signals warrants further examination. Genome-wide association studies (GWASs) and meta-analyses can shed light on the regions of the genome that tend to associate with a clinical phenotype, quantitative trait, or biomarker; this is accomplished through tagging and association-testing of single nucleotide polymorphisms (SNPs) that vary within the population. Recently developed methods like linkage disequilibrium (LD) score regression (LDSC)^23^ and Heritability Estimation from Summary Statistics (HESS)^24^ allow for direct comparison of GWAS summary statistics for two different phenotypes for quantitative assessment of genetic correlation.

In the present study, we leveraged existing data to explore the genetic associations of a set of medical phenotypes that are enriched with immune and inflammatory processes; these included allergic conditions, classic autoimmune diseases, other inflammatory diseases, and vulnerability to infectious disease. We sought to cross-correlate the genetic associations of these phenotypes with the associations obtained from studies of a set of psychiatric and behavioral phenotypes. We hypothesized that some phenotype-pairs with evidence for increased clinical comorbidity might also share similarities in their genome-wide association profile, which would be reflected in our analyses as significant positive correlations. Additionally, in light of literature suggesting shared genetic risk among some immune and inflammatory disorders, we assessed genetic correlations within this set of phenotypes using the LDSC method; these findings are reported within the Supplementary Materials. Genetic correlations within the set of psychiatric phenotypes have been reported previously^23,25,26^ and are not examined in the present study.

## Materials and Methods

### Literature Search

We searched the published literature (Pubmed, SCOPUS), data repositories (dbGaP and immunobase.org), and the downloads page of the Psychiatric Genomics Consortium (PGC) website (https://www.med.unc.edu/pgc/downloads) to identify phenotypes with potentially usable GWAS and GWAS meta-analysis summary statistics. For studies identified in the published literature, we contacted corresponding authors to request summary statistics. In order to facilitate cross-study comparison, we utilized studies that reported samples of European ancestry, broadly defined to include Central, Southern and Eastern Europe, Scandinavia, and Western Russia. Our initial search yielded a large number of datasets reflecting a wide-range of behavioral and immune-related phenotypes (Supplementary Table 1); the set of phenotypes ultimately retained for final analyses was selected based on criteria described below. When multiple studies were identified for a given phenotype, we pursued the studies with the largest effective sample sizes and ultimately used the available study with the largest heritability z-score. In several instances, data from the largest existing studies could not be shared or reflected a mixed-ancestry meta-analysis; in these cases, we deferred to the next largest European-ancestry study. We chose to retain datasets with an effective sample size greater than 5000 individuals and with estimated SNP heritability z-score > 3, in keeping with previous recommendations.^23^ This filter resulted in the exclusion of many relevant immune-related phenotypes, including eosinophilic esophagitis,^27^ granulomatosis with polyangiitis,^28^ IgA nephropathy,^29^ HIV-related neurocognitive phenotypes,^30^ morning cortisol levels,^31^ myeloid leukemias,^32^ psoriatic arthritis,^33^ sarcoidosis,^34^ and systemic sclerosis.^35^ This also resulted in exclusion of several psychiatric and behavior phenotypes, including adolescent alcohol abuse,^36^ anxiety-spectrum disorders,^37^ borderline personality disorder,^38^ language impairment,^39^ personality domains (five factor model),^40^ post-traumatic stress disorder,^41^ and reading disability.^42^ We also ultimately excluded data from studies of ethanol, opiate, and cocaine dependence,^43–45^ as genetic correlations involving these phenotypes were frequently outside the boundaries tolerated by the LDSC software, making them difficult to interpret; this may have been related to the ordinal-ranked phenotypes used in the GWASs. Finally, while relationships between tobacco-smoking behavior and other psychiatric phenotypes have been examined previously,^23,25^ we chose to retain smoking data in order to assess relationships with a more complete set of immune-related phenotypes. The full list of phenotypes identified in the search and retained for analyses is shown in Supplementary Table 1, along with identification of the study cohorts and consortia that generated these data, full citations of the respective publications, and indications of sample size, information regarding genomic inflation, and estimated SNP heritability.

**Figure 1.**
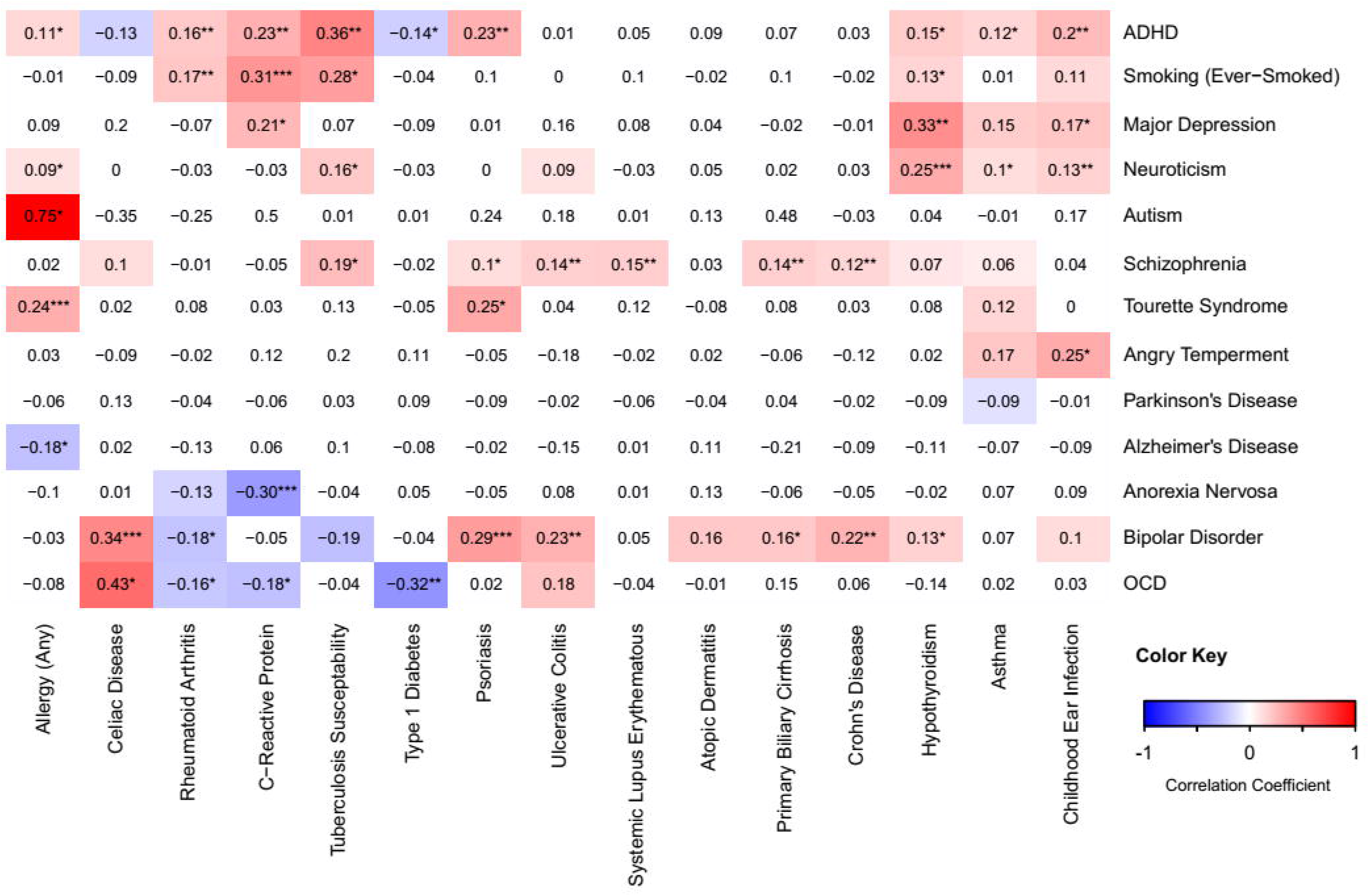
A heatmap depicting LDSC genome-wide genetic correlations between psychiatric and immune-related conditions such that red reflects more positive correlation coefficients while blue reflects more negative coefficients. Correlation coefficients are provided within each cell, with full details provided in Supplementary Table 2. Correlations reaching trend-level significance (0.05 < uncorrected *p* < 0.10) are depicted as colored panels, while relationships surpassing uncorrected *p* < 0.05 are additionally denoted with *, and relationships surpassing BH-*p* < 0.05 (for the total number of tests depicted in the figure) are denoted with ^**^. The rows and columns of the heatmap are hierarchically clustered based on correlation coefficients. Abbreviations: Attention deficit-hyperactivity disorder (ADHD), obsessive-compulsive disorder (OCD).

### GWAS Phenotypes Retained for Genetic Correlation

For our psychiatric and behavior-related phenotypes, we ultimately retained GWAS summary data reflecting studies of Alzheimer’s disease, ^46^ angry temperament ^47^, anorexia nervosa,^48^ attention deficit-hyperactivity disorder (ADHD),^49^ autism,^50^ bipolar disorder (BD),^51,52^ cigarette smoking (ever-smoked status),^53^ major depressive disorder,^54^ trait neuroticism,^55^ obsessive-compulsive disorder (OCD),^56^ Parkinson’s disease,^57^ schizophrenia (SZ),^58^ and Tourette Syndrome (personal communication from PGC Working Group). Collectively, these phenotypes were treated as a set. For phenotypes that are known or suspected to involve alterations to immune cells and/or inflammatory signaling, we ultimately retained GWAS data reflecting allergy (any, self-reported),^57,59^ asthma (self-reported),^57^ atopic dermatitis,^60^ childhood ear infection (self-reported),^57^ celiac disease,^61^ serum C-reactive protein (CRP),^62^ Crohn’s disease (CD),^63,64^ hypothyroidism (self-reported),^57^ primary biliary cirrhosis (PBC),^65^ psoriasis,^66^ rheumatoid arthritis,^67^ systemic lupus erythematosus (SLE),^68^ susceptibility to pulmonary tuberculosis,^69^ type 1 diabetes,^70^ and ulcerative colitis (UC).^71^ These phenotypes were treated as a set in subsequent analyses.

### Data Pre-Processing and Analysis

Our primary analyses were performed using the LDSC software (https://github.com/bulik/ldsc).^23^ Briefly, this set of tools can be used with existing GWAS summary data in order to distinguish polygenicity from confounding caused by uncontrolled population stratification or cryptic relatedness among samples,^72^ to estimate the heritability of a given phenotype,^23^ and to estimate the genetic correlation between two phenotypes based on two separate or related sets of summary statistics.^23^ In the latter application, the minimal requirements for each set of summary statistics include columns of data indicating SNP ID, the identities of reference and non-reference alleles, association *p*-value, effect size, test statistic (*e.g.,* odds ratio, regression β, or Z-score), and sample size (per SNP or for all SNPs). For each pair of phenotypes, this tool compares the strength and direction of association signal at each locus while correcting for the correlation that would be expected based on genetic linkage alone, and it provides an estimate of the genetic correlation between phenotypes. This method relies on adjustment for the linkage between SNPs (*i.e.,* covariance caused by genomic proximity); for our analyses, we used the set of LD scores provided by the software’s creators, based on the 1000 Genomes Project’s European sample (file = eur_w_ld_chr, URL = https://data.broadinstitute.org/alkesgroup/LDSCORE). Because minor allele frequencies (MAFs) and imputation quality scores were not available for all the obtained sets of GWAS results, we filtered the GWAS results to retain only SNPs that were included within the HapMap3 panel and had a MAF > 5 % within the 1000 Genomes Project Phase 3 European samples;^23^ this decision resulted in the exclusion of a sizable proportion of SNPs, but ensured equitable treatment of all datasets. The extended major histocompatibility complex (MHC) region contains high amounts of long-range LD, making it challenging to accurately map association signals in this region. For this reason, and following the work of others,^23,25^ we excluded this region from our analyses (chromosome 6, base positions 25x10^6^ to 35x10^6^). Additional SNP quality control (QC) routines followed those implemented by the GWAS authors and the defaults employed with the LDSC *munge_sumstats.py* function; this function checks alleles to ensure that the supplied alleles match those in the HapMap3 reference panel. For each dataset, we estimated the phenotype’s heritability. The results of this analysis, along with features of each GWAS dataset (sample size, number of QC-positive SNPs, genomic inflation factor, *etc.*), are shown for all phenotypes in Supplementary Table 1. All phenotypes with sample size > 5000 and estimated SNP heritability z-score > 3 were retained for correlation analysis (indicated in Supplementary Table 1 in green highlight). Pair-wise genetic correlations were assessed between retained phenotypes based on the intersection of QC-positive SNPs, and heatmaps were constructed to depict these relationships. For correlation coefficients returned within the bounds of the LDSC software, *p*-values were corrected using the Benjamini-Hochberg (BH) method for the total number of unique tests depicted in each correlation matrix. Within the main text, we describe only correlations that survived family-wise multiple-test correction. Correlations are reported as the coefficient ± standard error. For phenotype-pairs showing statistically significant genetic correlations, we re-evaluated the genetic correlations and estimated heritability using the HESS method (https://github.com/huwenboshi/hess).^24^

### Characterization of Genetically Correlated Loci and Associated Genes

For psychiatric-immune phenotype-pairs showing significant genetic correlations after BH correction for multiple testing, we used the HESS software to estimate partitioned heritability and genetic correlations based on smaller LD-based segments of the genome (average size = 1.5 Mb). We report the number and identity of genomic partitions (based on HG19 reference) displaying significant local genetic correlations and apply correction for the total number of partitions (≈1694, after MHC removal). Because presently available methods are poorly suited for fine-mapping the loci mediating a genetic correlation, we prioritized reporting correlated loci that also contain genome-wide significant associations for the relevant phenotypes (*i.e.*, associations with p < 5x10^−8^; subsequently called GW hits). We report GW hits contained within the present datasets, but also cross-reference these findings with those contained in immunobase.org, in order to identify loci associated with multiple immune-related disorders. We report the HGNC symbols for candidate genes proposed to mediate those associations. The full list of genes contained within each correlated loci is provided in Supplementary Table 3. Additionally, we used HESS to examine patterns of local genetic correlation in relationship to GWAS hits to make inferences about putative causal directionality between the phenotype-pairs. For all HESS analyses, we used the 1000 Genomes Project Phase 3 European reference panel and the LD-independent genome partitions recommended by the software developers.^73^ Following the developers’ practices, we assumed no sample overlap for comparisons of data generated by different consortia.^24^

## Results

### Genome-Wide Correlations between Psychiatric and Immune-Inflammatory Phenotypes

All pair-wise LDSC genetic correlations between psychiatric and immune-related phenotypes are depicted in Figure 1. Notably, twenty-one correlations survived BH correction for multiple testing (denoted with **) and 6 survived a more stringent Bonferroni correction (denoted with ***). Full results for these analyses are provided in Supplementary Table 2. Significant positive relationships were observed between ADHD and each of: CRP (*rg* = 0.23 ± 0.06, *p* = 2.0x10^−4^), childhood ear infections (*rg* = 0.20 ± 0.05, *p* = 2.0x10^−4^), psoriasis (*rg* = 0.23 ± 0.07, *p* = 1.0x10^−3^), rheumatoid arthritis (*rg* = 0.16 ± 0.05, *p* = 9.0x10^−4^), and tuberculosis susceptibility (*rg* = 0.36 ± 0.11, *p* = 1.6x10^−3^). Anorexia nervosa showed a negative genetic correlation with CRP (*rg* = −0.30 ± 0.08, *p* = 1.0x10^−4^). BD was positively correlated with each of: celiac disease (*rg* = 0.31 ± 0.09, *p* = 4.0x10^−4^), CD (*rg* = 0.21 ± 0.05, *p* = 3.7x10^−^ 5), psoriasis (*rg* = 0.25 ± 0.08, *p* = 3.8x10^−3^), and UC (*rg* = 0.23 ± 0.06, *p* = 2.0x10^−4^). Major depressive disorder was positively correlated with hypothyroidism (0.33 ± 0.09, *p* = 5.0x10^−4^). Similarly, neuroticism was positively correlated with hypothyroidism (*rg* = 0.25 ± 0.06, *p* = 7.2x10^−5^), in addition to childhood ear infection (*rg* = 0.13 ± 0.04, *p* = 8.0x10^−4^). OCD was negatively correlated with type 1 diabetes (*rg* = −0.32 ± 0.11, *p* = 5.4x10^−3^). Smoking behavior was positively correlated with CRP (*rg* = 0.31 ± 0.07, *p* = 3.6x10^−5^) and with rheumatoid arthritis (*rg* = 0.17 ± 0.05, *p* = 2.3x10^−3^). SZ showed positive genetic correlations with CD (*rg* = 0.12 ± 0.03, *p* = 2.0x10^−4^), PBC (*rg* = 0.14 ± 0.05, *p* = 2.0 x 10^−3^), SLE (*rg* = 0.15 ± 0.04, *p* = 2.0x10^−4^), and UC (*rg* = 0.14 ± 0.04, *p* = 2.0x10^−4^). Finally, we observed a positive genetic correlation between Tourette syndrome and allergy (*rg* = 0.24 ± 0.06, uncorrected *p* = 2.7x10^−5^). Additionally, several large-magnitude correlations attained a nominal threshold of statistical significance (*e.g.,* autism-allergy and OCD-celiac); these correlations tended to have a higher standard error and were generated using relatively smaller GWAS sample sizes. As such, they may be more likely to reflect false positives and should be regarded with appropriate skepticism.

For phenotypes involved in correlations that survived multiple test correction, estimated SNP heritability is shown in Table 2. For these phenotypes, we reassessed SNP heritability and the magnitude of genome-wide genetic correlations using the HESS method (Tables 1 and 2). Correlation coefficients were not correlated between the two methods (pearson *r* = 0.25, *p* = 0.25; Supplementary Figure 1) and the absolute value of the difference was negatively related to sample size (*r* = −0.45, *p* = 0.035; Supplementary Figure 2), which is consistent with the software developer’s guidelines.^24^ LDSC-based correlations among the immune-related phenotypes are reported in the Supplementary Text and Supplementary Table 5.

**Table 1:** Significant Genome-Wide Psychiatric-Immune Genetic Correlations. This table displays psychiatric-immune phenotype-pairs showing genome-wide genetic correlation with the linkage disequilibrium score regression (LDSC) method after correction for the total number of genetic correlations depicted in Figure 1 using the Benjamini-Hochberg (BH) method. We also report the genome-wide correlation estimates produced by the heritability estimation from summary statistics (*p*-HESS) method. Abbreviations: attention deficit-hyperactivity disorder (ADHD), bipolar disorder (BD), C-reactive protein (CRP), Crohn’s disease (CD), obsessive compulsive disorder (OCD), primary biliary cirrhosis (PBC), schizophrenia (SZ), systemic lupus erythematosus (SLE), ulcerative colitis (UC).

| Psychiatric Phenotype | Immune-Related Phenotype | LDSC Correlation ± Error, Uncorrected p-Value | HESS Correlation ± Error |
| --- | --- | --- | --- |
| ADHD | CRP | $0.23 \pm 0.06, p = 2.0 \times 10^{-4}$ | $0.21 \pm 0.04$ |
| ADHD | Childhood Ear Infection | $0.20 \pm 0.05, p = 2.0 \times 10^{-4}$ | $0.14 \pm 0.03$ |
| ADHD | Psoriasis | $0.23 \pm 0.07, p = 1.0 \times 10^{-3}$ | $1.99 \pm 0.20$ |
| ADHD | Rheumatoid Arthritis | $0.16 \pm 0.05, p = 9.0 \times 10^{-4}$ | $0.29 \pm 0.04$ |
| ADHD | Tuberculosis Susceptibility | $0.36 \pm 0.11, p = 1.6 \times 10^{-3}$ | $0.87 \pm 0.25$ |
| Anorexia Nervosa | CRP | $-0.30 \pm 0.08, p = 1.0 \times 10^{-4}$ | $-0.53 \pm 0.12$ |
| BD | Celiac Disease | $0.34 \pm 0.08, p = 4.5 \times 10^{-5}$ | $1.91 \pm 0.12$ |
| BD | CD | $0.22 \pm 0.06, p = 5.0 \times 10^{-4}$ | $1.31 \pm 0.07$ |
| BD | Psoriasis | $0.29 \pm 0.07, p = 2.7 \times 10^{-5}$ | $5.76 \pm 0.58$ |
| BD | UC | $0.23 \pm 0.07, p = 1.5 \times 10^{-3}$ | $1.59 \pm 0.08$ |
| Cigarettes (Ever-Smoked) | CRP | $0.31 \pm 0.07, p = 3.6 \times 10^{-5}$ | $2.24 \pm 0.73$ |
| Cigarettes (Ever-Smoked) | Rheumatoid Arthritis | $0.17 \pm 0.05, p = 2.3 \times 10^{-3}$ | $0.51 \pm 0.22$ |
| Major Depression | Hypothyroidism | $0.33 \pm 0.09, p = 5.0 \times 10^{-4}$ | $0.45 \pm 0.09$ |
| Neuroticism | Childhood Ear Infection | $0.13 \pm 0.04, p = 8.0 \times 10^{-4}$ | $0.06 \pm 0.01$ |
| Neuroticism | Hypothyroidism | $0.25 \pm 0.06, p = 7.2 \times 10^{-5}$ | $0.03 \pm 0.01$ |
| OCD | Type 1 Diabetes | $-0.32 \pm 0.11, p = 5.4 \times 10^{-3}$ | $0.98 \pm 0.18$ |
| SZ | CD | $0.12 \pm 0.03, p = 2.0 \times 10^{-4}$ | $0.31 \pm 0.03$ |
| SZ | PBC | $0.14 \pm 0.05, p = 2.0 \times 10^{-3}$ | $0.88 \pm 0.05$ |
| SZ | SLE | $0.15 \pm 0.05, p = 1.2 \times 10^{-3}$ | $0.12 \pm 0.02$ |
| SZ | UC | $0.14 \pm 0.04, p = 2.0 \times 10^{-4}$ | $0.56 \pm 0.03$ |
| Tourette Syndrome | Allergy (Any) | $0.24 \pm 0.06, p = 2.7 \times 10^{-5}$ | $0.29 \pm 0.07$ |

**Table 2:** Sample Characteristics for Phenotypes Involved in Significant Correlations. This table displays phenotype names, data sources, and estimated SNP heritability using the linkage disequilibrium score regression (LDSC) and heritability estimation from summary statistics (HESS) methods, as well as the GWAS sample size and number of SNPs surviving quality control. Full publication references, consortia names, links to web resources, and additional details on the original studies are provided in Supplementary Table I. GWAS N denoted with * indicates the median N for all SNPs. Abbreviations: Attention deficit-hyperactivity disorder (ADHD), bipolar disorder (BD), C-reactive protein (CRP), Crohn’s disease (CD), obsessive-compulsive disorder (OCD), primary biliary cirrhosis (PBC), Psychiatric Genomics Consortium (PGC), quality control (QC), single nucleotide polymorphism (SNP), schizophrenia (SZ), systemic lupus erythematosus (SLE), ulcerative colitis (UC).

| Phenotype | Data Source | Estimated Genome-Wide SNP Heritability ± Error (LDSC / HESS) | GWAS N | QC-Positive SNPs (MHC Excluded) |
| --- | --- | --- | --- | --- |
| ADHD | Demontis *et al.* (2017) | $0.24 \pm 0.02 / 0.26 \pm 0.02$ | 53,293 | 1,004,958 |
| Allergy (Any, Self-Report) | The 23andMe Research Team | $0.08 \pm 0.01 / 0.15 \pm 0.01$ | 181,000 | 1,060,611 |
| Anorexia Nervosa | Duncan *et al.* (2017) | $0.26 \pm 0.04 / 0.09 \pm 0.04$ | 14,477 | 1,054,719 |
| BP | Hou *et al.*, (2016) | $0.20 \pm 0.02 / 0.14 \pm 0.02$ | 40,225 | 1,052,397 |
| Childhood Ear Infection | The 23andMe Research Team | $0.07 \pm 0.01 / 0.10 \pm 0.01$ | 122,000 | 1,060,612 |
| Celiac Disease | Dubois *et al.*, (2010) | $0.30 \pm 0.05 / 0.13 \pm 0.04$ | 15,283 | 271,764 |
| Cigarettes (Ever-Smoked) | Tobacco and Genetics Consortium | $0.07 \pm 0.01 / 0.01 \pm 0.02$ | 74,035 | 963,355 |
| CD | Liu *et al.* (2015) | $0.47 \pm 0.06 / 0.33 \pm 0.03$ | 21,389 | 1,062,075 |
| CRP | Dehghan *et al.* (2011) | $0.13 \pm 0.02 / 0.11 \pm 0.02$ | 66,185 | 965,855 |
| Hypothyroidism (Self-Report) | The 23andMe Research Team | $0.05 \pm 0.01 / 0.08 \pm 0.01$ | 135,000 | 1,060,612 |
| Major Depression | PGC Depression Working Group | $0.14 \pm 0.03 / 0.07 \pm 0.04$ | 18,759 | 967,534 |
| Neuroticism | Social Science Genetics Consortium | $0.09 \pm 0.01 / 0.44 \pm 0.01$ | 168,105 | 1,053,712 |
| OCD | PGC OCD/TS Working Group | $0.29 \pm 0.05 / 0.09 \pm 0.04$ | 10,215* | 1,054,746 |
| PBC | Cordell *et al.*, (2015) | $0.37 \pm 0.06 / 0.17 \pm 0.04$ | 13,239 | 940,715 |
| Psoriasis | Tsoi *et al.*, (2015) | $0.82 \pm 0.13 / 0.09 \pm 0.04$ | 5,116* | 1,037,355 |
| Rheumatoid Arthritis | Okada *et al.*, (2014) | $0.14 \pm 0.02 / 0.10 \pm 0.01$ | 58,284 | 1,051,805 |
| SZ | PGC Schizophrenia Working Group | $0.47 \pm 0.02 / 0.62 \pm 0.01$ | 77,096 | 1,061,529 |
| SLE | Bentham *et al.*, (2015) | $0.27 \pm 0.05 / 0.27 \pm 0.03$ | 23,210 | 1,056,783 |
| Tourette Syndrome | PGC OCD/TS Working Group | $0.35 \pm 0.04 / 0.08 \pm 0.05$ | 13,341* | 1,041,689 |
| Tuberculosis Susceptibility | Curtis *et al.*, (2015) | $0.18 \pm 0.05 / 0.02 \pm 0.05$ | 11,936 | 819,917 |
| Type 1 Diabetes | Bradfield *et al.*, (2011) | $0.18 \pm 0.03 / 0.15 \pm 0.03$ | 26,890 | 854,164 |
| UC | Liu *et al.*, (2015) | $0.25 \pm 0.03 / 0.23 \pm 0.03$ | 27,432 | 1,062,094 |

### Characterization of Loci Contributing to Psychiatric-Immune Genetic Correlations

For psychiatric-immune phenotype-pairs that demonstrated a significant genome-wide correlation with the LDSC method (*i.e.,* those in Table 1), we used the HESS software to examine the genetic correlation within the ∼1694 partitioned genomic loci. The number of correlated loci before and after BH multiple test correction are depicted in Table 3; detailed results for these analyses, including local heritability, correlation strength, and the lists of gene symbols within each loci are provided in Supplementary Table 3. Only SZ displayed robust local genetic correlations with immune-related phenotypes, including thirty-two loci with CD, 37 loci with PBC, 20 loci with SLE, and 8 with UC (Table 3, depicted in Figure 2). Upon closer examination of the loci implicated between SZ and CD, we noticed that five of these loci contained GW hits, including one locus on chromosome 4q24 (4:100678360-103221356; highlighted green in Figure 2) that contained GW hits for both SZ and CD within the present data, and with 4 other autoimmune diseases (immunobase.org); these signals are near autoimmunity candidate genes *NFKB1* and *MANBA*, as well as proposed SZ candidate gene *SLC39A8*, among others contained within the locus (see Supplementary Table 3). The locus on 10p12.3 (10:18725659-18816236, highlighted green) contains a GW hit for SZ attributed to calcium channel gene *CACNB2*. Another locus mediating a significantly correlated locus on 12q12 (12:39227169-40816185, highlighted green) contains a GW hit for CD attributed to *LRRK2*. When examining the loci implicated between SZ and PBC, we observed 3 harboring GW hits for the former and 3 harboring signals for the latter, including loci within 3p24.3 (3:16282442-17891118, highlighted orange) containing *PLCL2* and within 11q23.3 (11:117747110-119215476, highlighted orange), containing candidate genes *CXCR5, DDX6*, and *TREH*. Among the loci implicated between SZ and SLE, we observed two harboring GW hits for the former and 3 harboring hits for the latter. One such locus within 1q21 (1:148361253-151538881, highlighted yellow) contains a SZ association signal localizing near candidate gene *APH1A.* Another locus within 1q23 (1:159913048-162346721, highlighted yellow) contains a GW hit for SLE, as well as several other autoimmune diseases, associated with candidate gene *FCGR2A*. Similarly, a locus within 22q11.21 (22:19912358-22357325, highlighted yellow) containing multi-disease association signal is associated with *MAPK1* and *UBE2L3*. Among the loci implicated between SZ and UC, one within 11q13.1 (11:63804569-65898631) harbored GW hits for multiple autoimmune disorders.

**Figure 2.**
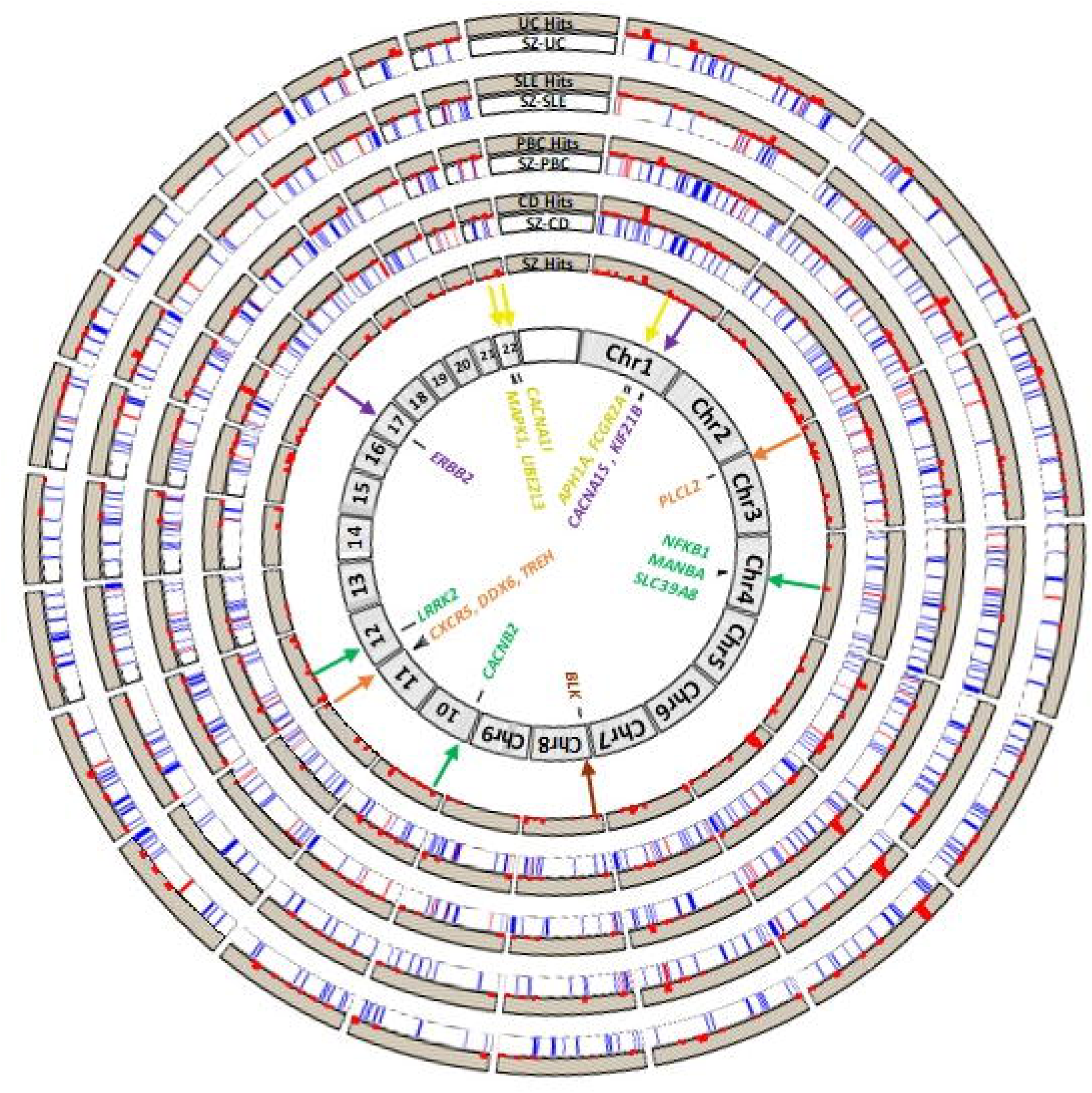
This figure depicts the HESS local genetic correlation data with respect to the genome and previously reported genome-wide association signals for respective disorders. A model genome using HG19 coordinates is depicted in grey. Moving outward from the center of the plot, the first track containing a red histogram depicts loci significantly associated with SZ (GWAS *p* < 5x10^−8^), with larger peaks indicating more significance (plotted as -log(*p*-value)). The second track (labeled SZ-CD) depicts regions of genetic correlation between SZ and CD, such that blue reflects uncorrected *p* < 0.05 and red reflects BH corrected *p* < 0.05. The next track (labeled CD Hits) contains a histogram depicting CD GWAS signal as described previously. The next pair of tracks depict genetic correlations for SZ-PBC and PBC GWAS signal, respectively. The third pair of tracks depicts this information for SZ-SLE (with SLE GWAS signal). The fourth pair of tracks depicts this information for SZ-UC and UC GWAS signal, respectively. In the center of the plot, we identify several GWAS candidate genes using colored text and arrows to indicate the pertinent locus; colored text and arrows are used to indicate the relevant phenotype-pairs, such that green = SZ-CD, orange = SZ-PBC, yellow = SZ-SLE, brown = SZ-PBC/SLE, and purple = SZ/BD-CD/PBC/UC.

**Table 3:**
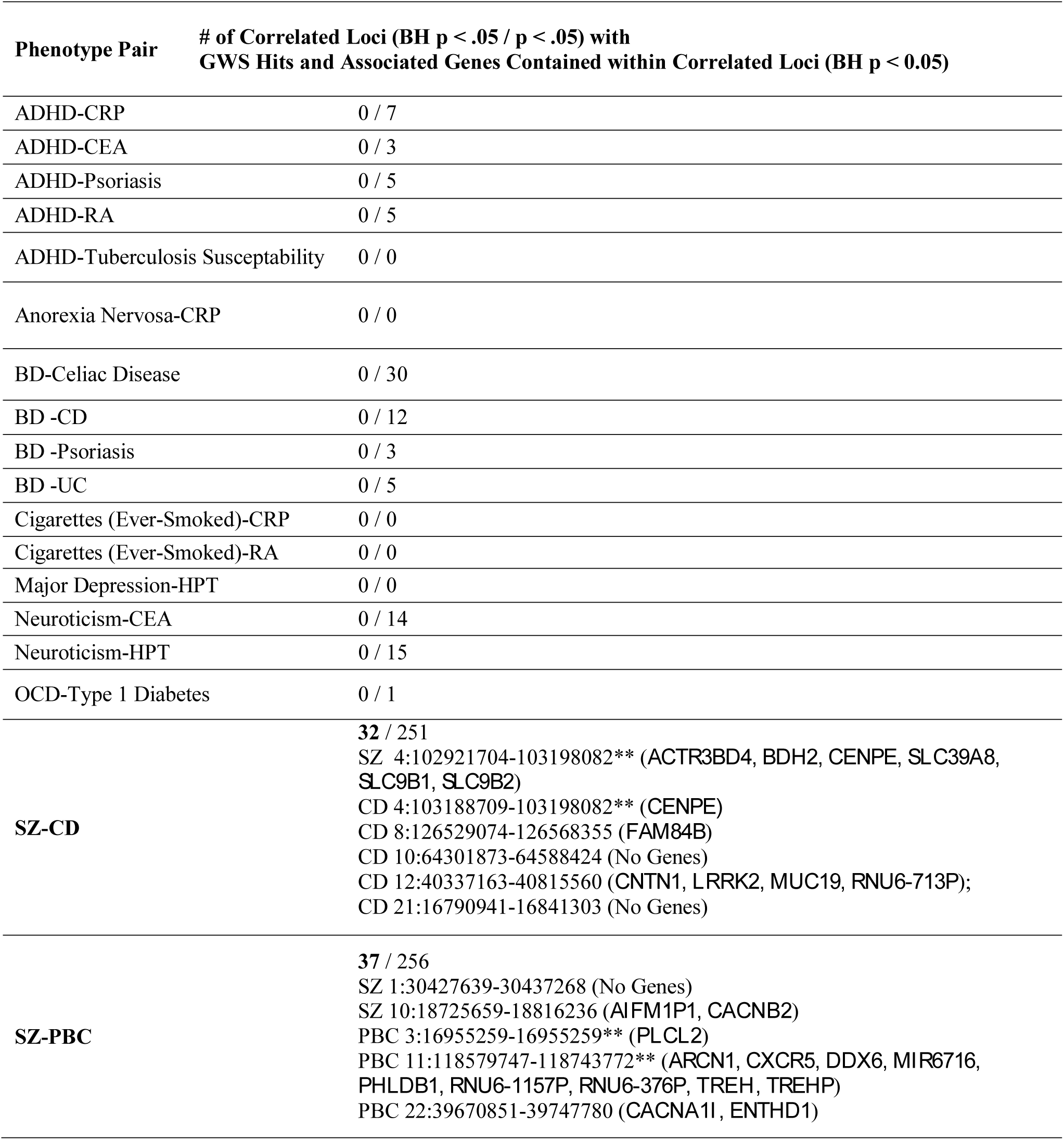
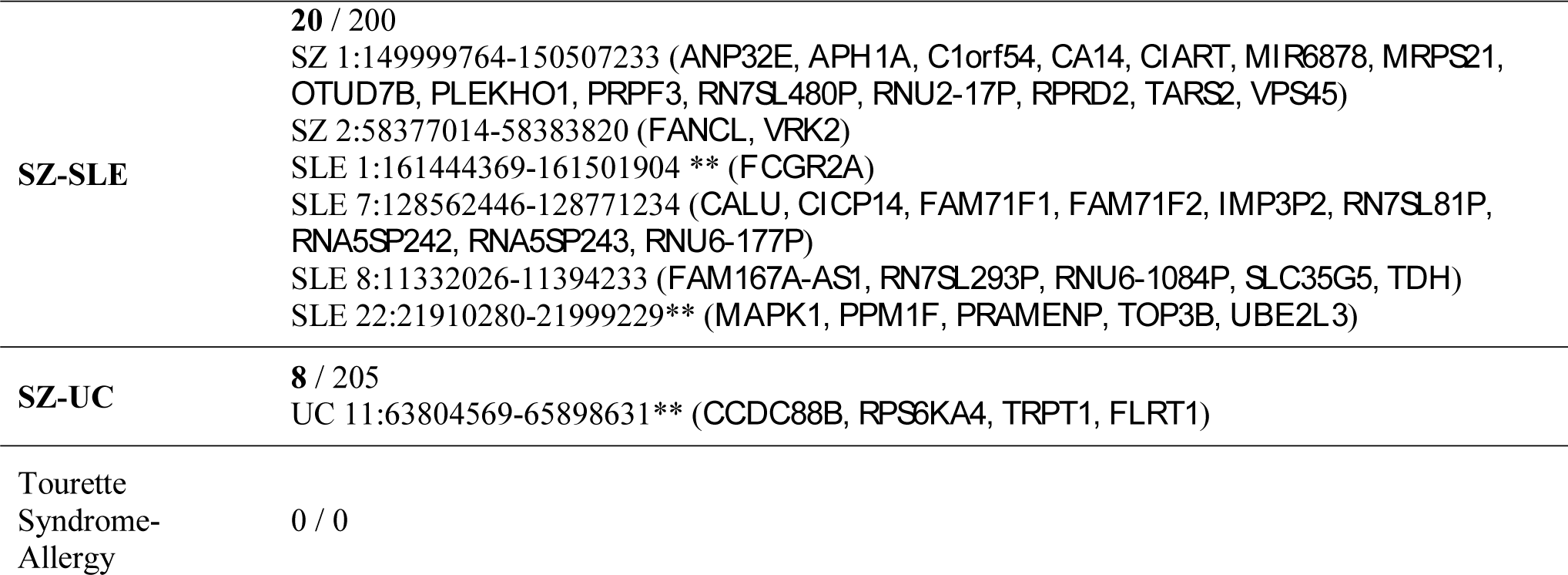
Significant Local Genetic Correlations Based on HESS Analysis. This table summarizes findings of local genetic correlation analysis, including the number of significantly correlated loci before and after Benjamini-Hochberg (BH) correction for multiple testing (**shown in bold**). Loci that showed robust correlations were interrogated for co-localization with significant genome-wide associations (GWS hits, with p < 5x10^−8^). The chromosomal coordinates containing GWS signal are provided, along with associated genes. Proposed candidate genes are highlighted with **bold text.** Abbreviations: Attention deficit-hyperactivity disorder (ADHD), Benjamini-Hochberg (BH), bipolar disorder (BD), childhood ear infection (CEA), Crohn’s disease (CD), hypothyroidism (HPT), comparison (NC-H), obsessive compulsive disorder (OCD), primary biliary cirrhosis (PBC), rheumatoid arthritis (RA), schizophrenia (SZ), systemic lupus erythematosus (SLE), ulcerative colitis (UC).

We also sought to examine whether the specific loci might be implicated across multiple psychiatric-immune disorder pairs (Figure 2). An analysis limited to only those surviving BH correction for multiple testing yielded only two loci shared by multiple disease pairs. The first locus (within 3p24.3; 3:21643707-22204244) was identified in correlations of SZ with PBC and with CD; it contained no GWS hits and two genes of unclear consequence *ZNF385D* and *ZNF385D-AS2*. The second locus within 8p32.1 (8:11278998-13491775, highlighted brown) was identified in correlations of SZ with PBC and with SLE; this locus contained numerous genes and is adjacent to a GWS hit for SLE associated with candidate gene *BLK*. When we broadened the scope to examine all loci implicated in nominally significant correlations (uncorrected *p* < 0.05), we find several that are common to multiple psychiatric-immune disorder pairs (Table 4). The most widely implicated locus was shared among the 5 pairs of psychotic and inflammatory bowel disorders (within 17q12; 17:36809344-38877404, highlighted purple) and contains a GW hit for BD ascribed to candidate gene *ERBB2*. There were another eight loci that were implicated in four disorder pairs. Among these, one located within 1q32.1 (1:200137649-201589975, highlighted purple) contains GW hits for multiple autoimmune disorders (including celiac disorder, CD, multiple sclerosis, and UC) and is near candidate genes *CACNA1S* and *KIF21B*. The full list of loci implicated across multiple disorder pairs is available in Supplementary Table 3. The results of the HESS analysis of putative causal directionality (Table 5) indicated that local genetic correlations were stronger in the loci containing GW hits for SZ (*rg* ≈ 0.41 ± 0.12) as compared with those containing hits for the paired autoimmune diseases (*rg* ≈ 0.17 ± 0.13).

**Table 4:** Loci Implicated Across Multiple Phenotype-Pairs at Uncorrected *p* < 0.05. This table depicts the loci that showed significant (uncorrected *p* < 0.05) correlations across multiple pairs of phenotypes. **Bold font** denotes phenotype-pairs for which the locus survived BH multiple test correction. The ** symbol denotes loci at which multiple autoimmune disorders show an association reaching genome-wide significance (per immunobase.org). **Bold font** is also used to indicate proposed candidate genes. Abbreviations: Bipolar disorder (BD), Crohn’s disease (CD), genome-wide significance (GWS) defined as *p* < 5x10^−8^, hypothyroidism (HPT), primary biliary cirrhosis (PBC), systemic lupus erythematosus (SLE), ulcerative colitis (UC).

| Locus | # of Pairs | Phenotype Pairs | GWS Associations and Nearby Genes |
| --- | --- | --- | --- |
| 17:36809344-38877404 | 5 | BD-CD, BD-UC, SZ-CD, SZ-PBC, SZ-UC | BP 17: 37839493-37893484 (***ERBB2***) / CD 17:37912377-38064876 / SLE 17: 38007190-38007319 / PBC 17:37912377-38080865 / UC 17: 37903731-38089717 (*RNU6-489P*, *TBC1D3C*, *TBC1D3D*, *TBC1D3K*, *TBC1D3L*) |
| 1:200137649-201589975 | 4 | BD-CD, BD-UC, **SZ-PBC**, SZ-UC | CD 1:200599616-201069559** / UC 1: 200864267-201024059** (*C1orf106*, *CACNA1S*, *GPR25*, ***KIF21B***) |
| 2:69139564-70755198 | 4 | BD-CD, SZ-CD, SZ-SLE, SZ-UC | None |
| 3:38356116-40221298 | 4 | Neuroticism-HPT, SZ-PBC, SZ-SLE, SZ-UC | None |
| 6:17386405-19207487 | 4 | SZ-CD, SZ-PBC, SZ-SLE, SZ-UC | None |
| 8:11278998-13491775 | 4 | Neuroticism-HPT, SZ-CD, **SZ-PBC**, **SZ-SLE** | Neuroticism 8:11281273-11895516 / SLE 8:11426026-11546260** (**BLK**, *C8orf49*, *CTSB*, *FAM167A*, *FAM167A-AS1*, *FDFT1*, *GATA4*, *LINC00208*, *MTMR9*, *NEIL2*, *RN7SL293P*, *RNU6-1084P*, *SLC35G5*, *SUB1P1*, *TDH*) |
| 8:9640787-10463197 | 4 | Neuroticism-HPT, SZ-PBC, SZ-SLE, SZ-UC | Neuroticism 8:9793601-10459000 (*LINC00599*, *MIR124-1*, *MSRA*) |
| 11:27020461-28481593 | 4 | SZ-CD, **SZ-PBC**, SZ-SLE, SZ-UC | None |
| 22:19912358-22357325 | 4 | SZ-CD, SZ-PBC, **SZ-SLE**, SZ-UC | CD 22:21916166-21985094** / SLE 22:21910280-21999229** (*CCDC116*, **MAPK1**, *RIMBP3*, *UBE2L3*, *YDJC*) |

**Table 5:** HESS Analysis of Putative Causal Directionality. Depicts the results of HESS analysis of putative causal directionality. Within this analysis, local genetic correlations are examined within loci containing GWS associations for each phenotype. The phenotype for which GWS loci produce the larger local correlations suggests that genetic liability for this phenotype may contribute to genetic risk for the other, especially when the correlation error bounds of the second phenotype overlap with zero. When both phenotypes show correlations overlapping with zero, no directionality is supported. Abbreviations: Crohn’s disease (CD), genome-wide significance (GWS) defined as p < 5x10^−8^, primary biliary cirrhosis (PBC), systemic lupus erythematosus (SLE), ulcerative colitis (UC).

| **Phenotype 1, Phenotype 2** | **Local Genetic Correlation ± Error at Loci Reaching GWS Only for Phenotype 1** | **Local Genetic Correlation ± Error at Loci Reaching GWS Only for Phenotype 2** | **Suggested Direction** |
| --- | --- | --- | --- |
| SZ-CD | $0.37 \pm 0.09$ | $0.11 \pm 0.08$ | SZ → CD |
| SZ-PBCs | $0.58 \pm 0.18$ | $0.26 \pm 0.17$ | SZ → PBC |
| SZ-SLE | $0.26 \pm 0.13$ | $0.16 \pm 0.16$ | SZ → SLE |
| SZ-UC | $0.43 \pm 0.09$ | $0.16 \pm 0.10$ | SZ → UC |

## Discussion

In contrast to previous studies examining large sets of medical, anthropomorphic, metabolic, and behavioral phenotypes,^23–26,74^ the present study performed a focused comparison of psychiatric and immune-related phenotypes using two methods to estimate genetic correlation from summary statistics. We used updated versions of psychiatric GWASs^49–51,56^ and compiled a more comprehensive set of immune-related phenotypes, while simultaneously reducing the burden imposed by multiple testing. Additionally, this analysis reflects the first application of the LDSC and HESS method for some of these phenotype-pairs. We identified several genome-wide correlations that were robust to multiple testing. Furthermore, we used the HESS method to validate genome-wide correlations and to conduct a quantitative analysis that localizes correlations to regions of the genome. We prioritized the reporting of findings based on co-localization with GW hits. As such, this study provides a quantitative map of genetic relationships between psychiatric and immune-related disorders and serves, along with previous work,^75^ as a starting point for identifying and characterizing potentially pleiotropic loci.

Prominent among the LDSC genome-wide significant findings was a cluster of modest positive correlations involving BD (*rg*s ranging 0.25 to 0.33) and SZ (*rg*s ranging 0.12 to 0.15) in conjunction with immune-related disorders involving the gastrointestinal tract (*i.e.,* CD, PBC, UC). These findings are consistent with available epidemiological evidence indicating that the presence of one set of disorders portends increased risk for a diagnosis from the other class of disorders, though the causality and temporality of these relationships is not clearly established.^76–82^ Positive genetic inter-correlations among these phenotypes are also consistent with recent work demonstrating that the positive correlation between BD and SZ are significantly mediated by both CNS and immunologic tissues.^83^ Our local genetic correlation analyses were inadequately powered to detect loci relevant to most of the psychiatric-immune disorder pairs, including BD. However, comparisons with SZ yielded 97 loci that were robust to multiple test correction, 18 of which also were shown to harbor GW hits in previous studies. In several instances, these GW hits localize near genes with functions that are pleiotropic and relevant to both brain and immune system phenotypes. For example, we identified a SZ-CD correlated locus at 4q24 (4:100678360-103221356) that contained GW hits for both SZ (putatively attributed to *SLC39A8*) and several autoimmune diseases (putatively attributed to *NFKB1* and *MANBA*); others have proposed that associations at this locus may exert pleiotropic effects on a wide range of phenotypes (additionally including body mass index, serum levels of manganese, N-terminal pro-B-type natriuretic peptide, and HDL-cholesterol) through a functional variant found in European populations affecting the *SLC39A8* cation transporter.^84,85^ A locus within 11q23.3 (11:117747110-119215476) was significantly correlated between SZ and PBC and harbors a region of GW hits for multiple autoimmune disorders attributed to *PLCL2*, a catalytically inactive phospholipase-like protein thought to influence intracellular signaling, calcium homeostasis, and GABA-ergic receptor trafficking in immune and neuronal cell types, among others.^86–88^ A *de-novo* missense mutation affecting this gene was identified in an exome sequencing study of SZ affected individuals, though no replication appears to have been reported.^89^ Similarly, a correlated locus within 22q13.1 (22:39307894-40545797, highlighted yellow in Figure 2) contains GW hits for PBC, which overlaps with voltage gated calcium channel gene *CACNA1I*; this gene has been implicated by both GWAS and rare-variant studies of SZ.^58,90^ Another correlated locus within 11q23 (11:118579747-118743772) contained GW hits for multiple autoimmune disorders and is suspected to exert pleiotropic effects through several genes, whose functions include repression of aberrant interferon signaling (*DDX6*),^91^ chemokine signaling between T-helper and B-cells (*CXCR5*),^92,93^ and enzymatic break down of microbial disaccharides (*TREH).*^94^ Notably, functional genomic studies have identified *DDX6* as a gene that is perturbed during neuronal differentiation of samples derived from individuals with schizophrenia,^95^ and as a peripheral blood marker of cerebrospinal fluid serotonin metabolite levels,^96^ supporting its relevance to psychiatric phenotypes.

We also examined loci that showed a nominal genetic correlation across multiple disorder pairs, and found these loci also harbored GW hits for respective phenotypes. The locus at 17q12 shared among multiple disorders contains a GW hit for BD (17:36809344-38877404) ascribed to candidate gene *ERBB2*.^52^ This gene and its relatives encode receptor tyrosine kinases that interact with a family of growth factors called neuregulins to regulate the assembly of neural circuitry, myelination, neurotransmission and synaptic plasticity. A large body of evidence implicates both ligands and receptors from these families as susceptibility genes for SZ and BD.^97^ Notably, *ERBB2* overlaps with GW hits for multiple autoimmune disorders, though these have been attributed to different genes in the region. Another locus at 1q32.1 (1:200137649-201589975) contains GW hits for multiple autoimmune disorders (including celiac disease, CD, multiple sclerosis, and UC) and is near candidate genes *C1orf106, CACNA1S, GPR25*, and *KIF21B*. Genetic disruptions of voltage-gated calcium channels, including *CACNA1S,* are well-established susceptibility factors in psychiatric and neurological disorders.^98,99^ *KIF21B* encodes a neuronal motor protein implicated in GABA_A_ receptor trafficking,^100^ in addition to having a suspected role in regulating inflammatory signaling in several lymphocyte subtypes.^101^

While it is tempting to speculate about these observations, we must acknowledge limitations and caveats of the present approach. Current methods for assessing genetic correlations are not well-suited for fine-mapping shared liability across disorders; other methods are better suited for this task, including extensions of GWAS that model multiple phenotypes simultaneously.^22,55,102,103^ With respect to local genetic correlations, we have prioritized reporting of loci that co-localize with GW hits. However, this implies that the presence of the GW hit is contributing to the observed correlation, which we have not demonstrated presently. As such, our discussion of potentially pleiotropic loci and candidate genes should be considered anecdotal at this time. One indirect approach to assessing the role of GW hits in a local genetic correlation might be to re-estimate the local correlation after the removal of the smaller region of GW signal from the original datasets. When we conducted this analysis for the SZ-CD pair, we found that the number of significant loci (BH *p* < 0.05) was reduced from 32 to 8, suggesting that GW hits likely play an important role in many of the local genetic correlations. Future studies will be able to combine larger GWAS sample sizes with new methods aimed at stratifying genetic correlations by biological annotations (*e.g.,* tissue type or signaling pathways) in order to more precisely define the parts of the genome that mediate a genetic correlation.^83^

Several methods have now been used to examine quantitative SNP-based genetic relationships between psychiatric and immune-related phenotypes, including restricted maximum likelihood (REML) co-heritability, polygenic risk scores, genetic analysis incorporating pleiotropy and annotations, and other permutation-based methods.^22,104–106^ Different approaches rest on unique assumptions, test different sets of hypotheses, and appear prone to generating sometimes conflicting results. Using several approaches that were not dependent on the directionality of a given SNP’s effect, Wang and colleagues concluded that many (24 of 35) pairs of psychiatric and immune-related phenotypes shared a statistically significant proportion of risk-associated loci; among these findings was a significant genetic overlap between BD (as well as SZ) and UC.^22^ However, many of the other relationships identified in that study were not significant in the present study. Another recent study demonstrated that polygenic risk scores reflecting additive risk for several autoimmune diseases can explain a small proportion of variance in SZ case-control status, yet the genome-wide significant SNPs from the autoimmune GWASs were not over-represented among SZ’s genome-wide significant hits when permutation-based analysis was performed.^105^ The apparent disagreement between different approaches for assessing shared genetic liability thus underscores the value of examining the consensus across studies and methods.^105^

The LDSC approach featured here attempts to quantitate similarities and differences in association signals across the entire genome. Some of our phenotype-pairs have been examined previously using genome-wide assessment methods, yielding apparently contradictory findings.^25,26,72^ For example, a previous study implementing a REML-based approach did not find significant SNP-based co-heritabilities between CD and the major psychiatric phenotypes.^106^ Additionally, the first study implementing the LDSC method found no significant correlation (*rg* = 0.08 ± 0.08, uncorrected *p* = 0.33) between BD and UC;^23^ this study used a smaller dataset for BD (Sklar *et al*., 2011; *N* = 16,731) and a different version of the UC dataset (reported as Jostins *et al.,*2012; *N =* 27,432). A similar non-correlation is also reported in LD-Hub (http://ldsc.broadinstitute.org/), using what appears to be the same datasets, although referencing a related article (Liu *et al.*, 2015; *N* = 27,432). The analyses portrayed in our main text utilized a larger BD dataset (Hou *et al.*, N = 40,225), the same dataset for UC (Liu *et al.*, 2015; N = 27,432), and uniform criteria for SNP retention based on inclusion in the HapMap3 panel and MAF > 5 % within the 1000 Genomes Project Phase 3 European samples. In order to resolve apparent discrepancies, we obtained additional versions of the available data for BD, SZ, CD, and UC and pre-filtered under both inclusive (imputation INFO score > 0.9 or all SNPs, when INFO score unavailable) or exclusive criteria (MAF > 5 % within the 1000 Genomes Project Phase 3 European samples). We found that correlations between SZ and each of CD, PBC, and UC tended to be more positive and more significant (*i.e.,* reaching a BH-corrected threshold) when using the SZ data filtered at MAF > 5% (Supplementary Figure 3). A similar pattern held true for inclusive *vs*. exclusive pre-filtering for the BD dataset generated by Sklar *et al.*, but this was not the case for the larger Hou *et al.*, dataset. A side-by-side comparison of the effects of different pre-filtering decisions for the BD, SZ, CD, and UC datasets in relation to the other phenotypes is provided in Supplementary Figure 4. These observations indicate that decisions pertaining to SNP inclusion can have a considerable effect on the result of the LDSC analysis; this idea is further supported by the observation that stratified genetic correlation analyses based on MAF thresholds can produce different levels of statistical significance and opposite patterns of correlation directionality.^83^ Thus, our study suggests that genetic correlations between psychiatric and immune-related disorders may be more significant when analyses are restricted to common variation. Reassuringly, the developers of the HESS method use the same datasets examined presently, and also report positive genetic correlations between SZ and the inflammatory bowel disorders.^24^ The results of the HESS analysis of putative causal directionality indicate that the local genetic correlations are higher in loci occupied by SZ GW hits, as compared to the loci harboring hits for the paired autoimmune disorders.^24^ This pattern is consistent with the hypothesis that genetic liability toward SZ tends to impart a greater genetic risk for the corresponding paired disorder, rather than the opposite directional hypothesis. A related interpretation may be there is an unobserved intermediate phenotype (*e.g.,* a shared biological pathways/mechanism) that is pleiotropic for both measured phenotypes, but more strongly influences the SZ phenotype. This pattern of findings could also be caused by the presence of a confounding factor (*e.g.,* smoking, socioeconomic status) that portends risk for both phenotypes.^24^ Thus, we caution against over-interpretation of these findings. Extensions of Mendelian randomization methods to incorporate two GWAS samples using multi-allelic risk stratifying instruments will be better suited to address these hypotheses,^107^ especially as future GWASs provide well-powered genetic estimates of potentially relevant intermediate phenotypes (*e.g.,* brain structure morphometry, circulating immune cell phenotypes, and serum cytokine levels).^108–110^ Other limitations of the HESS method, including assumptions related to sample overlap and ancestry stratification, are discussed extensively by the method’s developers.^24^

Our study also identified many phenotype-pairs that demonstrated significant genome-wide correlations using the LDSC method, but for which HESS-based genome-wide and local genetic correlations could not be identified. This is unsurprising, given that the sample sizes for these phenotypes were generally below the recommended sample size for HESS analyses (*N* > 50,000).^24^ Nonetheless, some of these relationships are supported by evidence from clinical and epidemiological studies, and thus may warrant follow-up using larger sample sizes and alternative methods for assessing genetic relationships. For example, we observed a modest positive correlation between self-reported hypothyroidism and major depression (*rg* = 0.33 ± 0.09, *p* = 5.0x10^−4^), as well as trait neuroticism (*rg* =0.25 ± 0.06, *p* = 7.2x10^−5^). This could be consistent with two different sets of clinical observations. The first is that symptoms of depression are common in individuals with hypothyroidism, and that subclinical hypothyroidism could play a role in a subset of persons diagnosed with major depression; thus cross-contamination of GWAS samples could lead to a biased positive correlation. However, the second observation is that there is an increased incidence of major depression and depressive symptomatology in persons with autoimmune thyroiditis receiving hormone replacement therapy.^111,112^ It is worth noting that GWAS data for allergy, asthma, hypothyroidism, childhood ear infection, and Parkinson’s disease were obtained through 23andMe, Inc.. These data are based on self-report, and thus could be more susceptible to bias stemming from misdiagnosis or misreporting, though previous work supports their validity.^113^ None the less, the samples sizes are an order of magnitude larger than many other datasets, resulting in smaller standardized errors and better power for the detection of weak genetic correlations. It is yet unclear whether small magnitude genetic correlations like these might be clinically meaningful. The LDSC correlations observed presently were relatively weak magnitude (*rg*s ≈ 0.12 to 0.30) and of modest modest statistical significance (1x10^−5^ < uncorrected *p* < 5x10^−3^), when compared to the strongest genetic correlations observed within each group of datasets (*e.g.,* SZ-BD *rg* = 0.87 with *p* = 7.4x10^−94^; CD-UC *rg*= 0.71 with *p =* 3.5x10^−36^).

Several other significant genetic correlations are supported in the clinical and epidemiological literature. For example, we found a positive correlation between ADHD and rheumatoid arthritis (*rg* =0.16 ± 0.05, *p* = 9.0x10^−4^); this finds support in large registry-based studies indicating an increase in ADHD diagnosis in individuals with autoimmune disease,^114^ children with mother’s affected by autoimmune disease,^114^ and children of mothers with rheumatoid arthritis.^115^ Registry-based studies also provide support for increased incidence of ear infections (*rg* = 0.20 ± 0.05, *p* = 2.0x10^−4^) and psoriasis (*rg*= 0.23 ± 0.07, *p* = 1.0x10^−3^) among individuals with ADHD. ^114,116–118^ On the other hand, ADHD was positively correlated with CRP (*rg* = 0.23 ± 0.06, *p* = 2.0x10^−4^), though a relatively large epidemiological study finds no association in affected individuals.^119^ The negative correlation between anorexia nervosa and CRP (*rg* = −0.30 ± 0.08, *p* = 1.0x10-4) is borne out in a recent meta-analysis of relevant studies.^120^ Another negative correlation between OCD and type 1 diabetes (*rg* = −0.32 ± 0.11, *p* = 5.4x10^−3^) finds no support within a limited body of literature.^121^ However, the positive correlation between Tourette syndrome and allergy (*rg* = 0.24 ± 0.06, *p* = 2.7x10^−5^) is consistent with evidence of increased comorbidity between these phenotypes.^122,123^ There is a paucity of clinical studies directly assessing the relationship between SZ and PBC (*rg* = 0.14 ± 0.05, *p* = 2.0 x 10^−3^). On the other hand, the correlation between SZ and SLE (*rg* = 0.15 ± 0.04, *p* = 2.0x10^−4^) appears to be supported by both epidemiological evidence of increased comorbidity^124^ and the well-documented (although rare) phenomenon of CNS lupus presenting with SZ-like symptoms,^125^ which may contribute to misdiagnosis. Finally, positive correlations involving cigarette smoking behavior and CRP (*rg* = 0.31 ± 0.07, *p* = 3.6x10^−5^), as well as rheumatoid arthritis (*rg* = 0.17 ± 0.05, *p* = 2.3x10^−3^), are perhaps unsurprising given considerable evidence of elevated CRP in persons who smoke,^126^ and increased incidence of smoking behavior among individuals diagnosed with rheumatoid arthritis.^127^ These findings may indicate a need for more adequate statistical treatment of smoking behavior in GWAS studies.

The present study identified a number of intriguing and previously unreported genetic correlations, some of which appear to localize near established risk factors for complex disease. On the whole, these findings are consistent with the idea that similar signatures of common genetic variation may increase risk for both psychiatric and immune-related disorders. However, it is important to keep in mind that these findings do not necessarily imply causality or even shared genetic etiology. SNP-based genetic correlations could arise from a wide variety of underlying factors, including the possibility that the relationship between phenotypes is mediated by behavioral or cultural factors, or influenced by a heritable but unexamined underlying trait that confers risk to both phenotypes.^23,26^ Other factors that could contribute to genetic correlations include effects mediated by parental genotypes and their influence on parental behaviors that impact the offspring.^128^ Additionally, GWAS studies of psychiatric phenotypes typically do not screen affected cases on the presence of other medical conditions (and *vice-versa*), thus over-representation of a given phenotype in the sample of another phenotype could bias the data toward the detection of a genetic correlation. Finally, estimates of genetic similarities could be influenced by misdiagnosed cases.^129^ Other general limitations of this method (in comparison with other approaches) have been discussed previously elsewhere.^23,26^ In light of the exploratory nature of the present study, another critique pertains to the lack of clearly identified positive and negative control comparisons. Additionally, the clinical significance of weak or modest genetic correlations is yet unclear. Future work could shed light on this topic by comparing the strength of reported genetic correlations with estimates of effect size from epidemiological associations, in order to create an atlas of concordance and shed light on the sensitivity and specificity of these genetic methods. One final critique of this approach is that it falls short of identifying plausible genetic and biological mechanisms that mediate potentially pleiotropic loci. Future work incorporating expression quantitative trait loci, differentially expressed or methylated genes, or enriched ontological and functional terms may provide a clearer context for assessing biological similarities between phenotypes. Despite these limitations, the present study indicates that shared aspects of common genetic variation may underlie long-recognized epidemiological links between psychiatric and immune-related disorders and serves as a start point for the identification and characterization of potentially pleiotropic loci.

## Acknowledgements

The authors gratefully acknowledge the contributions of all the individuals (patients, families, research participants, clinicians and diagnosticians, research associates, and data analysts) and consortia whose efforts made possible the GWAS studies and meta-analyses featured in the present study. For most of the phenotypes examined in the present study, clinical and genetic data were collected across numerous sites, each with their own unique patients, staff, and funding sources. While we attempted to provide more thorough recognition of the required acknowledgments for each individual phenotype in our supplementary note, we realize that it is not possible to recognize every individual and funding mechanism that made these studies possible, and we apologize for this. We gratefully acknowledge 23andMe, Inc., its staff, and its customers who consented to participate in research. We also gratefully acknowledge the developers of the LDSC and HESS softwares. We gratefully acknowledge Susan Service for her assistance preparing and analyzing the data supporting the original association studies of neurocognitive impairment and dementia in HIV-affected adults.

## Funding Sources

The authors declare no conflicts of interest related to this study. Authors S.J.G., D.S.T., and J.L.H were supported by the U.S. National Institutes of Health [R01MH101519, R01AG054002]; the Sidney R. Baer, Jr. Foundation; NARSAD: The Brain & Behavior Research Foundation; and the Gerber Foundation [awarded to S.J.G.]. D.S.T. was also supported by Autism Speaks [Weatherstone Pre-doctoral Training Grant #9645]. R.M. was supported by the Vascular Dementia Research Foundation. A.J.L. was supported by the UCLA-AIDS Institute and the UCLA Center for AIDS Research [AI28697 awarded to Freimer & A.J.L.]; as well as NIH R03DA026099 [awarded to A.J.L.]. S.N. is a Wellcome Trust Senior Research Fellow in Basic Biomedical Science and is also supported by the NIHR Cambridge Biomedical Research Centre.

## Legends for Tables and Supplementary Tables

Table 1. This table displays psychiatric-immune phenotype-pairs showing genome-wide genetic correlation with the linkage disequilibrium score regression (LDSC) method after correction for the total number of genetic correlations depicted in Figure 1 using the Benjamini-Hochberg (BH) method. We also report the genome-wide correlation estimates produced by the heritability estimation from summary statistics (*p*-HESS) method. Abbreviations: attention deficit-hyperactivity disorder (ADHD), bipolar disorder (BD), C-reactive protein (CRP), Crohn’s disease (CD), obsessive compulsive disorder (OCD), primary biliary cirrhosis (PBC), schizophrenia (SZ), systemic lupus erythematosus (SLE), ulcerative colitis (UC).

Table 2. This table displays phenotype names, data sources, and estimated SNP heritability using the linkage disequilibrium score regression (LDSC) and heritability estimation from summary statistics (HESS) methods, as well as the GWAS sample size and number of SNPs surviving quality control. Full publication references, consortia names, links to web resources, and additional details on the original studies are provided in Supplementary Table I. GWAS N denoted with ^*^ indicates the median N for all SNPs. Abbreviations: Attention deficit-hyperactivity disorder (ADHD), bipolar disorder (BD), C-reactive protein (CRP), Crohn’s disease (CD), obsessive-compulsive disorder (OCD), primary biliary cirrhosis (PBC), Psychiatric Genomics Consortium (PGC), quality control (QC), single nucleotide polymorphism (SNP), schizophrenia (SZ), systemic lupus erythematosus (SLE), ulcerative colitis (UC).

Table 3. This table summarizes findings of local genetic correlation analysis, including the number of significantly correlated loci before and after Benjamini-Hochberg (BH) correction for multiple testing (**shown in bold**). Loci that showed robust correlations were interrogated for co-localization with significant genome-wide associations (GWS hits, with p < 5x10^−8^). The chromosomal coordinates containing GWS signal are provided, along with associated genes. Proposed candidate genes are highlighted with **bold text.** Abbreviations: Attention deficit-hyperactivity disorder (ADHD), Benjamini-Hochberg (BH), bipolar disorder (BD), childhood ear infection (CEA), Crohn’s disease (CD), hypothyroidism (HPT), comparison (NC-H), obsessive compulsive disorder (OCD), primary biliary cirrhosis (PBC), rheumatoid arthritis (RA), schizophrenia (SZ), systemic lupus erythematosus (SLE), ulcerative colitis (UC).

Table 4. This table depicts the loci that showed significant (uncorrected *p* < 0.05) correlations across multiple pairs of phenotypes. **Bold font** denotes phenotype-pairs for which the locus survived BH multiple test correction. The ^**^ symbol denotes loci at which multiple autoimmune disorders show an association reaching genome-wide significance (per immunobase.org). **Bold font** is also used to indicate proposed candidate genes. Abbreviations: Bipolar disorder (BD), Crohn’s disease (CD), genome-wide significance (GWS) defined as *p* < 5x10^−8^, hypothyroidism (HPT), primary biliary cirrhosis (PBC), systemic lupus erythematosus (SLE), ulcerative colitis (UC).

Table 5. Depicts the results of HESS analysis of putative causal directionality. Within this analysis, local genetic correlations are examined within loci containing GWS associations for each phenotype. The phenotype for which GWS loci produce the larger local correlations suggests that genetic liability for this phenotype may contribute to genetic risk for the other, especially when the correlation error bounds of the second phenotype overlap with zero. When both phenotypes show correlations overlapping with zero, no directionality is supported. Abbreviations: Crohn’s disease (CD), genome-wide significance (GWS) defined as *p* < 5x10^−8^, primary biliary cirrhosis (PBC), systemic lupus erythematosus (SLE), ulcerative colitis (UC).

Supplementary Table 1. GWAS Sample Information and Single Phenotype Statistics, MHC Excluded.

Supplementary Table 2. LDSC Psychiatric-Immune Correlations

Supplementary Table 3. HESS Local Genetic Correlations.

Supplementary Table 4. LDSC Immune-Immune Correlations

**Supplementary Figure 1.**
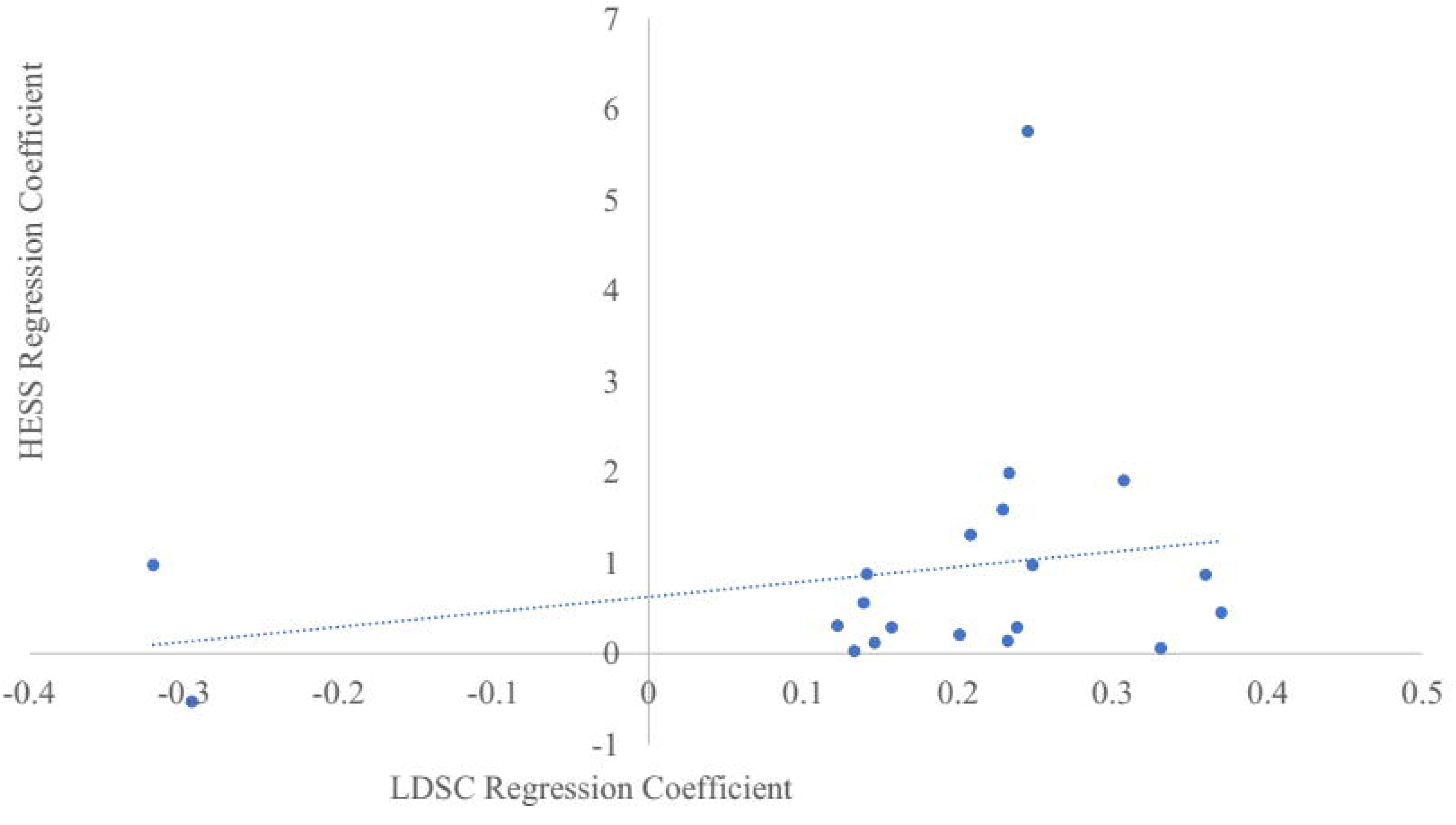
Depicts the relationship between LDSC and HESS genome-wide genetic correlation coefficients (pearson *r* = 0.25, *p* = 0.25).

**Supplementary Figure 2.**
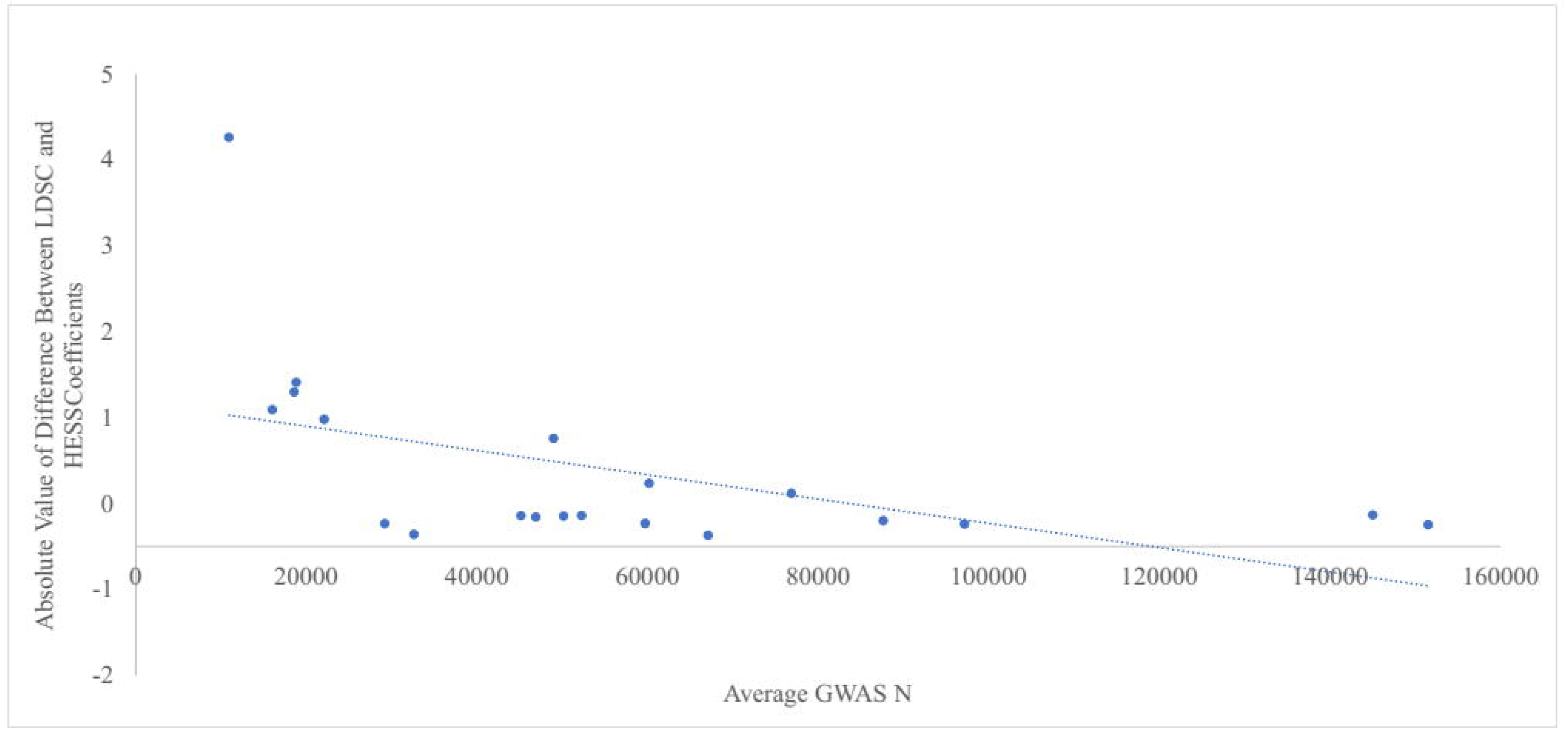
Depicts the absolute value of the difference between LDSC and HESS genome-wide genetic correlation coefficients and the average sample size of the two contributing GWAS studies (pearson *r* = −0.44, *p* = 0.035).

**Supplementary Figure 3.**
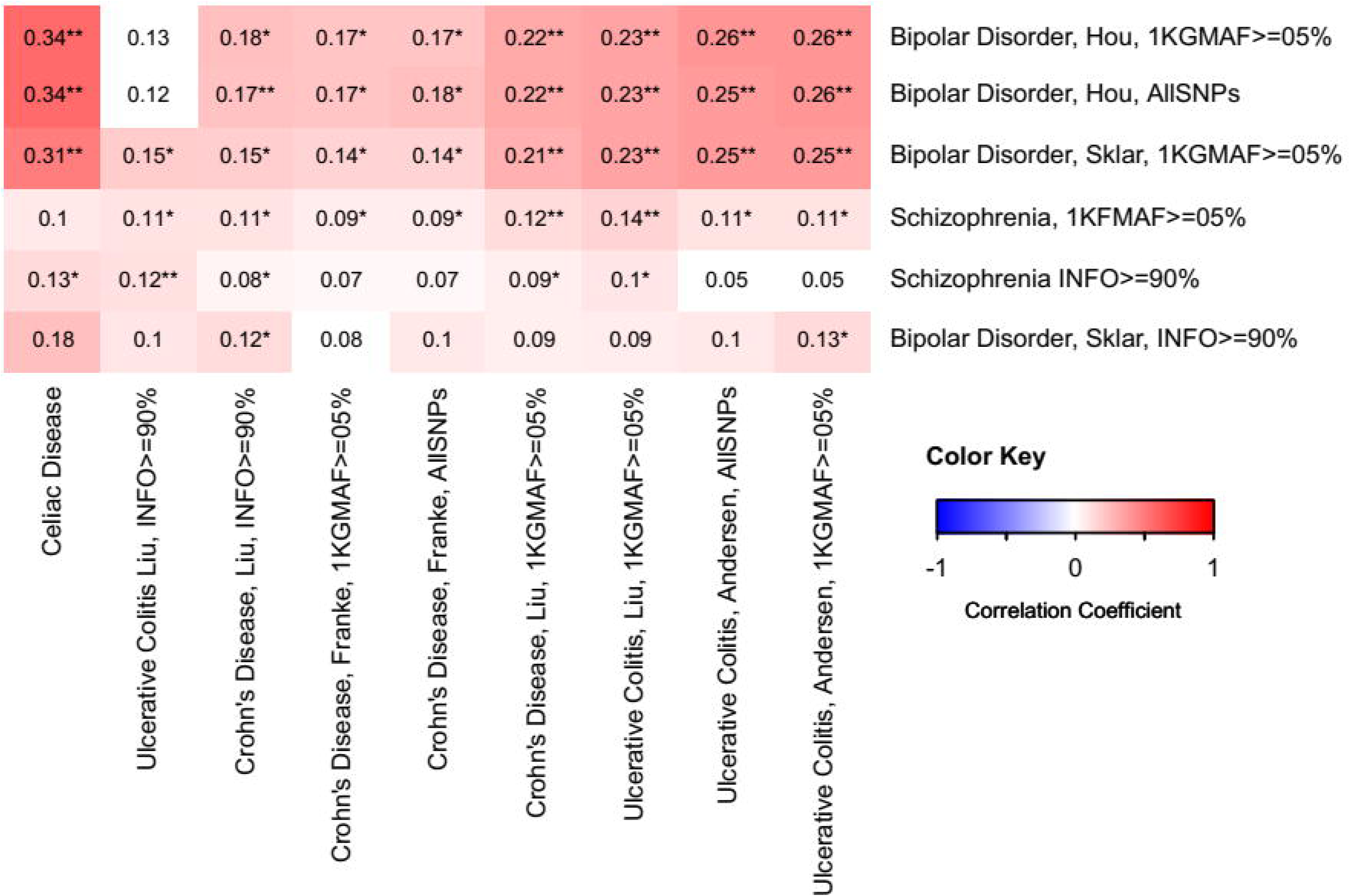
Depicts differences in LDSC-based genome-wide correlations based on dataset selection and pre-filtering decisions for a select set of phenotypes. Each dataset is coded with the GWAS first author’s name and the filtering threshold. 1KGMAF>=05% reflects retention of SNPs with minor allele frequencies > 5% within the thousand genomes phase 3 reference panel. INFO>=90% reflects retention of SNPs with imputation quality scores > 0.9. ALL SNPs indicates that no SNPs were filtered, because INFO score was not available for these data. Correlations reaching trend-level significance (0.05< uncorrected *p* < 0.10) are depicted as colored panels, while relationships surpassing uncorrected *p* < 0.05 are additionally denoted with ^*,^ and relationships surpassing BH-*p* < 0.05 (for the total number of tests depicted in Supplementary Figure 2) are denoted with ^**^.

**Supplementary Figure 4.**
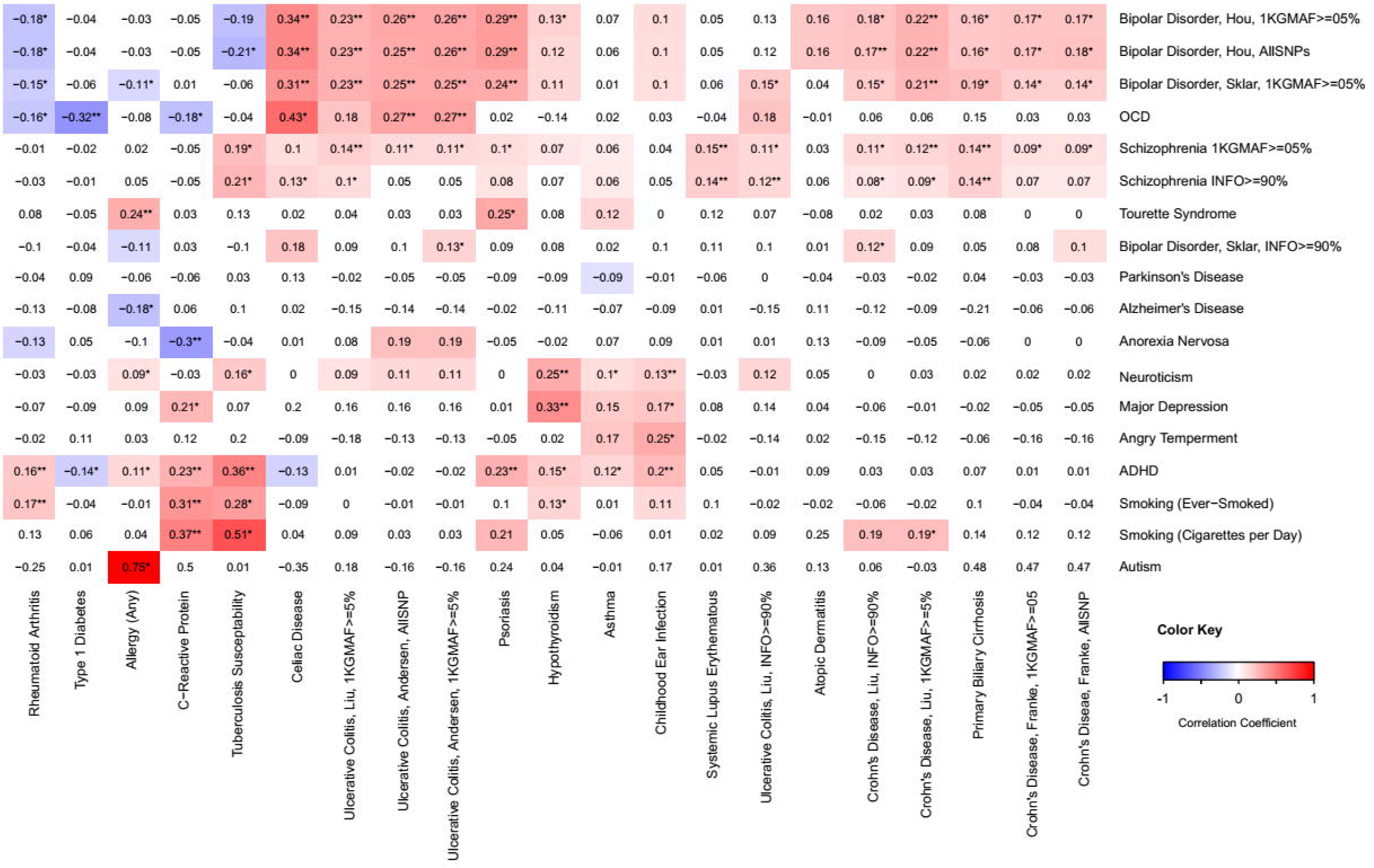
Depicts differences in LDSC-based genome-wide correlations based on dataset selection and pre-filtering decisions for a select set of phenotypes in relation to the larger set of phenotypes. Datasets that were differentially processed are indicated with first author’s name and the filtering threshold. 1KGMAF>=05% reflects retention of SNPs with minor allele frequencies > 5% within the thousand genomes phase 3 reference panel. INFO>=90% reflects retention of SNPs with imputation quality scores > 0.9. ALL SNPs indicates that no SNPs were filtered, because INFO score was not available for these data. Correlations reaching trend-level significance (0.05 < uncorrected *p* < 0.10) are depicted as colored panels, while relationships surpassing uncorrected *p* < 0.05 are additionally denoted with ^*,^ and relationships surpassing BH-*p* < 0.05 (for the total number of tests depicted in Supplementary Figure 2) are denoted with ^**^. Full results are provided in Supplementary Table 2.

**Supplementary Figure 5.**
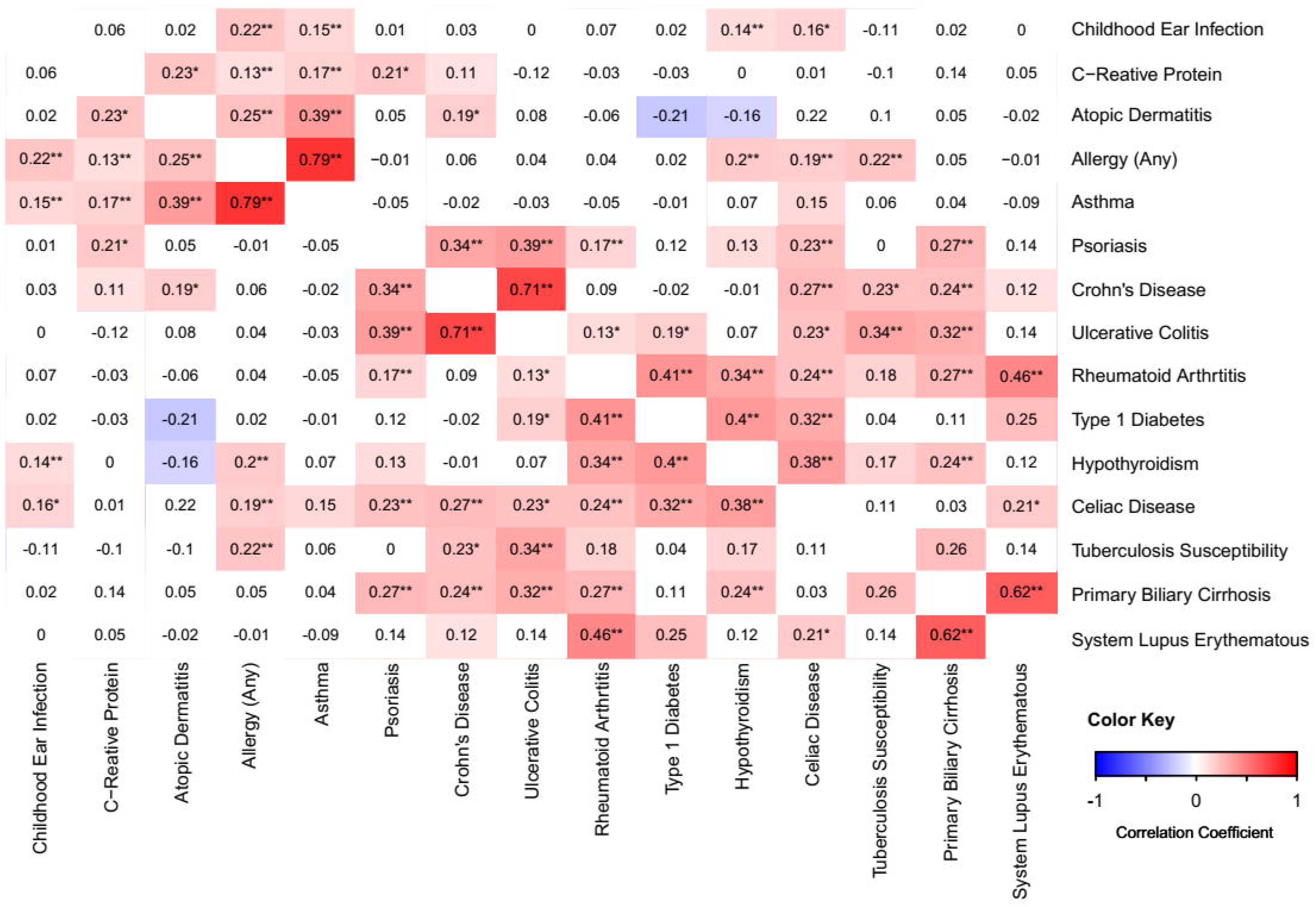
A heatmap depicting LDSC genome-wide genetic correlations between among immune-related conditions such that red reflects more positive correlation coefficients while blue reflects more negative coefficients. Correlation coefficients are provided within each cell, with full details provided in Supplementary Table 4. Correlations reaching trend-level significance (0.05 < uncorrected *p* < 0.10) are depicted as colored panels, while relationships surpassing uncorrected *p* < 0.05 are additionally denoted with ^*,^ and relationships surpassing BH-*p* < 0.05 (for the total number of tests depicted in the figure) are denoted with ^**^. The rows and columns of the heatmap are hierarchically clustered based on correlation coefficients.

## Manuscript Abbreviations

(ADHD): Attention deficit-hyperactivity disorder
(BH): Benjamini-Hochberg
(BD): bipolar disorder
(CRP): C-reactive protein
(CD): Crohn’s disease
(GWAS): genome-wide association study
(GW hits): genome-wide significant associations
(HESS): heritability estimation from summary statistics
(LD): linkage disequilibrium
(LDSC): linkage disequilibrium score regression
(MHC): major histocompatibility
(OCD): obsessive compulsive disorder
(PBC): primary biliary cirrhosis
(PGC): Psychiatric Genomics Consortium
(QC): quality control
(REML): restricted maximum likelihood
(SZ): schizophrenia
(SNP): single nucleotide polymorphism
(SLE): systemic lupus erythematosus
(UC): ulcerative colitis

